# In-planta Gene Targeting in Barley using Cas9, with and without Geminiviral Replicons

**DOI:** 10.1101/2021.02.04.429787

**Authors:** Tom Lawrenson, Alison Hinchliffe, Martha Clarke, Yvie Morgan, Wendy Harwood

## Abstract

Advances in the use of RNA-guided Cas9-based genome editing in plants have been rapid over the last few years. A desirable application of genome editing is gene targeting (GT), as it allows a wide range of precise modifications, however this remains inefficient especially in key crop species. Here we describe successful, heritable gene targeting in barley using an *in-planta* strategy but fail to achieve the same using a wheat dwarf virus replicon to increase copy number of the repair template. Without the replicon, we were able to delete 150bp of the coding sequence of our target gene whilst simultaneously fusing in-frame mCherry in its place. Starting from 14 original transgenic plants, two plants appeared to have the required gene targeting event. From one of these T0 plants, three independent gene targeting events were identified, two of which were heritable. When the replicon was included, 39 T0 plants were produced and shown to have high copy numbers of the repair template. However, none of the 17 lines screened in T1 gave rise to significant or heritable gene targeting events despite screening twice the number of plants in T1 compared to the non-replicon strategy. Investigation indicated that high copy numbers of repair template created by the replicon approach cause false positive PCR results which are indistinguishable at the sequence level to true GT events in junction PCR screens widely used in GT studies. In the successful non-replicon approach, heritable gene targeting events were obtained in T1 and subsequently the T-DNA was found to be linked to the targeted locus. Thus, physical proximity of target and donor sites may be a factor in successful gene targeting.

## Introduction

Genome editing has exploded in recent years due to advances in programmable nucleases which allow a double stranded DNA break to be created at a predefined locus. First on the scene were Zinc-finger nucleases (Kim et al., 1996) followed by transcription activator-like effector nucleases (TALENS) (Christian et al., 2010) and more recently clustered regularly interspaced short palindromic repeats (CRISPR) systems, especially the *Sp* Cas9 (Jinek et al., 2012) which was the first CRISPR nuclease reported to function in plants (Feng et al., 2013; Li et al., 2013; Nekrasov et al., 2013; Shan et al., 2013; Xie and Yang, 2013). Although insertion of exogenously supplied DNA into plant genomes has been possible for many years via *Agrobacterium*-mediated transformation or physical delivery, location was impossible to control precisely. Some success is reported inserting DNA in a precise manor by homologous recombination in rice without creating a double strand break (DSB) at the target site, although it was necessary to use a positive/negative selection system (Terada et al., 2002) which was later shown to produce no successful modifications in barley (Horvath et al., 2017). The value of creating a DSB at the target site to initiate DNA repair and facilitate insertion by homologous recombination was shown early on in plants with the non-programmable *I-SceI* meganuclease (Fauser et al., 2012), so it was a natural progression to repurpose Cas9 for precise insertional modifications. Many DSBs are repaired by non-homologous end joining (NHEJ) mechanisms which are error prone but shown to be capable of inserting an exogenously supplied DNA template at the break point in plants (Salomon and Puchta, 1998; Chilton and Que, 2003; Tzfira et al., 2003; Lee et al., 2019) although precision is likely to be compromised by indels created at the spliced junctions as well as issues controlling orientation, truncation and concatenation.

Targeted DSBs can be introduced very efficiently and with great precision into plant genomes using RNA-guided Cas9 and this has made it facile over recent years to produce gene knockouts by the introduction of indels and larger deletions due to the error prone nature of NHEJ. Many reports now exist describing single and multiple gene knockouts at efficiencies often approaching 100% although the precise nature of the edit is often not possible to predict. Lesions typically lead to a shift in reading frame and a premature stop codon. Base and prime editing technologies (Komor et al., 2016; Gaudelli et al., 2017; Anzalone et al., 2019) have partially addressed the precision issue allowing defined single base changes as well as short insertions and deletions, although larger precise changes such as adding an in-frame reporter fusion are unlikely to be possible in this way.

Gene targeting (GT) can be defined as the introduction of a precise predefined modification into a plant genome, either an insertion, deletion or replacement via the introduction of a supplied repair template using homology dependent recombination (HDR) and usually a DSB at the target site. By making available a repair template containing the required modification flanked by sequence homologous to each side of the DSB, a precise change can be introduced into the genome. This change can be either small, for example a single amino acid conversion (Budhagatapalli et al., 2015; Sun et al., 2016; Svitashev et al., 2016; Wolter et al., 2018; Danilo et al., 2019; Wolter and Puchta, 2019), or large such as the in-frame insertion of a reporter gene (Zhao et al., 2016; Wang et al., 2017; Miki et al., 2018).

Whilst GT is able to address both large and small precise modifications it is usually much harder to achieve than knock out and so researchers have sought ways in which rare events can be screened for easily and means to boost the frequency at which they occur. Early GT efforts in crops have focused on creating a precise change resulting in resistance to a herbicide or antibiotic which can then be used to select for resistant plants containing the desired GT event. *ALS* (acetolactate synthase) is a plant gene essential in the production of branched chain amino acids that is a target for inhibitors used as herbicides which has been extensively used in plants for GT experiments (Svitashev et al., 2015; Sun et al., 2016; Svitashev et al., 2016; Wolter et al., 2018; Danilo et al., 2019). Sometimes a visual marker has been used in the screen such as insertion of a 35s promoter upstream of *ANT1* leading to a purple phenotype (Cermak et al., 2015), or restoration of *gl1* leading to trichome production in *Arabidopsis* (Hahn et al., 2018). This approach however means that the modification is restricted to genes which allow such a selectable or visible phenotype, which many editing projects will not.

Many crop plants may only be transformed at efficiencies of a few percent or less, which, when combined with the low efficiency of GT makes regeneration of T0 gene targeted plants hugely labour intensive or just inconceivable. One way around this is to adopt an in-planta strategy whereby just a few primary transgenics containing the editing reagents are created, but the numbers required to retrieve the rare GT events are generated by the plants themselves through the normal process of flowering and seed production (Fauser et al., 2012; Schiml et al., 2014; Schiml et al., 2017). Each progeny plant may give rise to successful GT events, perhaps just as somatic sectors, but these can enter the germ line and prove to be heritable in subsequent generations. In this approach, all the editing reagents can be included on a single T-DNA with a selection cassette to allow transgenic production, a nuclease programmed to create a DSB at the target site and a repair template containing the desired modification with flanking sequence homologous to each side of the target site DSB. Recognition sequences for the nuclease can also be added to the ends of the repair template to allow cutting and its transfer to the target site (Schiml et al., 2014; Zhao et al., 2016). Here the screen can be based on the genotype rather than the phenotype, plants containing the required edits being detected by PCR for example. A widely adopted approach is to PCR screen using one primer within the modified region of the repair template and the second primer outside of the repair template in the sequence flanking the target site. In this way the PCR must cross the junction where the repair template stops, and the flanking genomic sequence begins.

It has been suggested that one major constraint on successful GT is the availability of repair template sequence at the correct time and in sufficient quantity for it to be incorporated as intended. In order to address this, Geminivirus replicons have been utilised (Baltes et al., 2014; Butler et al., 2016; Gil-Humanes et al., 2017; Wang et al., 2017; Dahan-Meir et al., 2018; Hahn et al., 2018; Vu et al., 2020), simultaneously pushing plant cells into more of an S phase-like state where HDR repair occurs more frequently, and by replicating to high copy numbers providing many copies of the repair template to the target site. Here the coat and movement protein section of the viral genome can be replaced by the repair template and then supplied in linear form to the plant on a T-DNA for *Agrobacterium*-mediated delivery. Once within the plant cell, the viral REP proteins are expressed leading to rolling circle replication and many copies of the repair template. Cas9 can also be delivered on the same T-DNA allowing simultaneous DSB at the target site and production of large quantities of the repair template. This approach has been most successful in tomato (Cermak et al., 2015; Dahan-Meir et al., 2018; Vu et al., 2020) but has also been described in wheat (Gil-Humanes et al., 2017), rice (Wang et al., 2017) and potato (Butler et al., 2016) although *Arabidopsis* appeared to be recalcitrant to any GT benefits (Hahn et al., 2018).

To date there have been two reports of GT in barley, one a transient single amino acid conversion of a GFP transgene to YFP (Budhagatapalli et al., 2015) and the second a stable modification of a non-functional *hptII* transgene to a functional form (Watanabe et al., 2016). The former was identified in 3 epidermal cells and the latter were one-sided GT events - one side of the repair was by HDR and the other by NHEJ. Our aim was to achieve heritable Cas9 GT in barley which would modify a locus of interest that was not a transgene and not chemically or visibly selectable, thus we chose to create a partial deletion of a native barley gene of interest, simultaneously fusing an in-frame reporter to the remaining part. To keep the number of transgenics required to a minimum and to potentially make the approach suitable for genotypes more recalcitrant to transformation, we used an in-planta strategy and attempted to increase efficiency by incorporating the repair template within a Geminivirus replicon. We present efficiencies using strategies with and without inclusion of the replicon.

## Materials and Methods

### PCR and Sanger sequencing of HORVU4Hr1G061310 target locus for indel detection

PCR was done using 30ng of genomic DNA as template, 400nM of primers F4 & R5 (supplementary file 8), Qiagen 2x PCR mastermix and a total reaction volume of 25μl being completed by water. After initial denaturation at 94°C for 3 minutes, 40 cycles of: 94°C for 30 seconds/58°C for 1 minute/72°C for 45 seconds were performed. A final extension of 72°C for 5 minutes was given. 1μl of the cleaned product (see Sanger sequencing PCR products) was then used in separate Sanger reactions with both F4 and R5 primers. ABI chromatograms were observed for indications of indels or larger deletions between guides A and B either as homozygous modifications or mixtures of alleles indicated by the sudden appearance of multiple peaks at the points where Cas9 is known to cut (3bp from PAM).

### Sanger sequencing of PCR products

PCR reactions were prepared for Sanger sequencing by adding 10 units of exonuclease 1 and 1unit of shrimp alkaline phosphatase to 10μl of PCR reaction before incubation at 37°C for 30 minutes followed by 80°C for 10 minutes to inactivate the enzymes. 1μl of the cleaned product was used as sequencing template where the amplicon was 1Kb or less in size and 2μl when over 1Kb. Sequencing reactions were in 10μl volumes with 100nM primer, 1.5μl Big Dye buffer and 1μl Big Dye 3.1, made up to 10μl with water. After a denaturation step of 96°C for 2 minutes, 35 cycles of: 96°C for 10 seconds/52°C for 15 seconds/60°C for 3 minutes were performed. Finally, reactions were held at 72°C for 1 minute, sent for commercial data extraction and return in the form of ABI files.

### qPCR assay for T-DNA (HptII)/& repair template (mCherry) copy number determination

Hydrolysis probe based quantitative real-time PCR (qPCR) was used to determine copy number of the T-DNA (*HptII*) and repair template (mCherry) in transgenic barley lines. The reaction compared the Cq values of an *HptII* amplicon to a single-copy barley gene *CO2* (*Constans-like*, AF490469) amplicon and the Cq values of an mCherry amplicon to a single-copy barley gene *CO2* (*Constans-like*, AF490469) amplicon within FAM/VIC duplexed assays (see supplementary file 8). The reactions used Thermo ABGene Absolute qPCR Rox Mix (Cat number AB1139) with the probes and primers at a final concentration of 200 nM (*HptII* and mCherry) and 100nM (*CO2*). The assay contained 5 μL DNA solution and was optimised for DNA concentrations of 1 to 10 ng/μL (5 to 50 ng DNA in the assay). PCRs were carried out as 25μl reactions in a Bio-Rad CFX96 machine (C1000 Touch). The detectors used were FAM-TAMRA and VIC-TAMRA. The PCR cycling conditions were 95 °C for 15 min (enzyme activation), 40 cycles of 95 °C for 15 s, and 60 °C for 60 s. Cq values were determined using CFX96 software (version 3.1), with Cq determination set to regression mode. Values obtained were used to calculate copy number according to published methods (Weng et al., 2004).

### PCR screening for GT

F1/R1 (left junction) and F2/R2 (right junction) primer sequences are given in supplementary file 8. Each left and right junction PCR reaction contained 30ng genomic DNA template, 2.5μl 10X buffer 1, 200μM dNTPs, 200nM primers, 0.625units amplitaq gold (Thermo fisher) and water to 25μl. Reactions were cycled: 95°C 10 minutes (enzyme activation), then 40 cycles of 95°C for 30 seconds/58°C for 30 seconds/72°C for 1 minute. Final extension was at 72°C for 5 minutes. Amplicons were sequenced with the following primers (supplementary file 8): F1/R1 amplicon: Seq1, Seq2, Seq3, Seq10. F2/R2 amplicon: Seq6, Seq7, Seq8, Seq9, Seq1. Primers for the less sensitive but fully diagnostic F1/R3 PCR are given in supplementary file 8. Each reaction contained 30ng genomic DNA template, 10ul 5X GoTaq buffer, 1.5mm MgCl_2_, 200nM primers, 200μM dNTPs, 5 units GoTaq DNA polymerase (Promega) and water to 50μl. Reactions were cycled at 94°C for 3 minutes then 40 cycles of 94°C for 30 seconds/58°C for 30 seconds/72°C for 2 minutes and 30 seconds, before final extension at 72°C for 5 minutes. F1/R3 amplicons were sequenced with the following primers (supplementary file 8): Seq1, Seq 2, Seq 3, Seq 4, Seq 5, Seq6, Seq7, Seq8, Seq10.

### gDNA prep and quantification by Qubit

Genomic DNA was prepared from leaves according to a published protocol (Edwards et al., 1991). Preps were quantified using the Qubit dsDNA HS Assay Kit (Thermo Fisher) according to the manufacturer’s instructions and diluted to a concentration of 30ng/μl.

### Barley transformation

Barley (*cv.* ‘Golden Promise’) was transformed by *Agrobacterium tumefaciens-* mediated transformation of immature embryos as described by Hinchliffe and Harwood (Hinchliffe and Harwood, 2019)

### Construct assembly

Constructs were assembled using previously described parts and methods (Lawrenson et al., 2015), except for the protospacers, repair template and WDV components. Protospacers, repair template, extended repair template and replicon sequences are given in supplementary file 8. Protospacer, repair template, extended repair template and replicon sequences were commercially synthesised as modules compatible with the parts and cloning methods previously described (Lawrenson et al., 2015). Sequence confirmed constructs A,B,C and D were transformed into *Agrobacterium* strain AGL1.

### Crossing

Barley was crossed according to a published protocol (Thomas et al., 2019).

### Chromosome walking

A published PCR based protocol (Wang et al., 2007) was used to determine sequences flanking the T-DNA borders. Primer sequences used are shown in supplementary file 8. SP2 products were cloned into pGEMT-Easy (Promega) and sequenced with M13 and M13R universal primers (Eurofins). pGEMT-Easy, left and right T-DNA border sequences were identified in the reads showing that the remaining barley sequence represented the sequence flanking the T-DNA in the barley genome.

## Results

### GT Construct Design

In our design strategy, high efficiency introduction of DSBs was considered important as the benefits of DSBs to GT have been previously reported (Fauser et al., 2012). Therefore, for our selected native barley target (HORVU4Hr1G061310), two protospacers were identified that gave good results and would allow for a strategy to delete around 150bp of the coding sequence of this single exon gene whilst simultaneously fusing in-frame mCherry in its place (figure 1.1a-c). Protospacer A was able to create indels, as detectable by Sanger sequencing of PCR amplicons covering the target site, in 9/18 (50%) of independent transgenic lines and protospacer B 16/18 (89%) of the same lines. To maximise the chance of success, we decided to incorporate both guides into our design as two DSBs at the target site might be better than one. In the repair template, homology to the target site was maximised by continuing the right and left homology arm sequences fully up to the Cas9 cuts sites i.e. 3bp from the native PAM. This allowed omission of the PAM on the left arm and the protospacer on the right arm of the repair template, preventing the Cas9 from cutting within it both before and after GT (figure 1.1b). Target sequences (full protospacer and PAM) were included in the flanks of the repair template (figure 1.2) to allow cutting and facilitate its incorporation into the target site. The repair template was added to the construct containing Cas9 and the guide A & B cassettes to arrive at construct A (figure 1.2a) which was similar in architecture to a previous example shown to enable GT in *Arabidopsis* (Schiml et al., 2014). The predicted GT event is shown in figure 1.1c.

**Figure 1.1.**
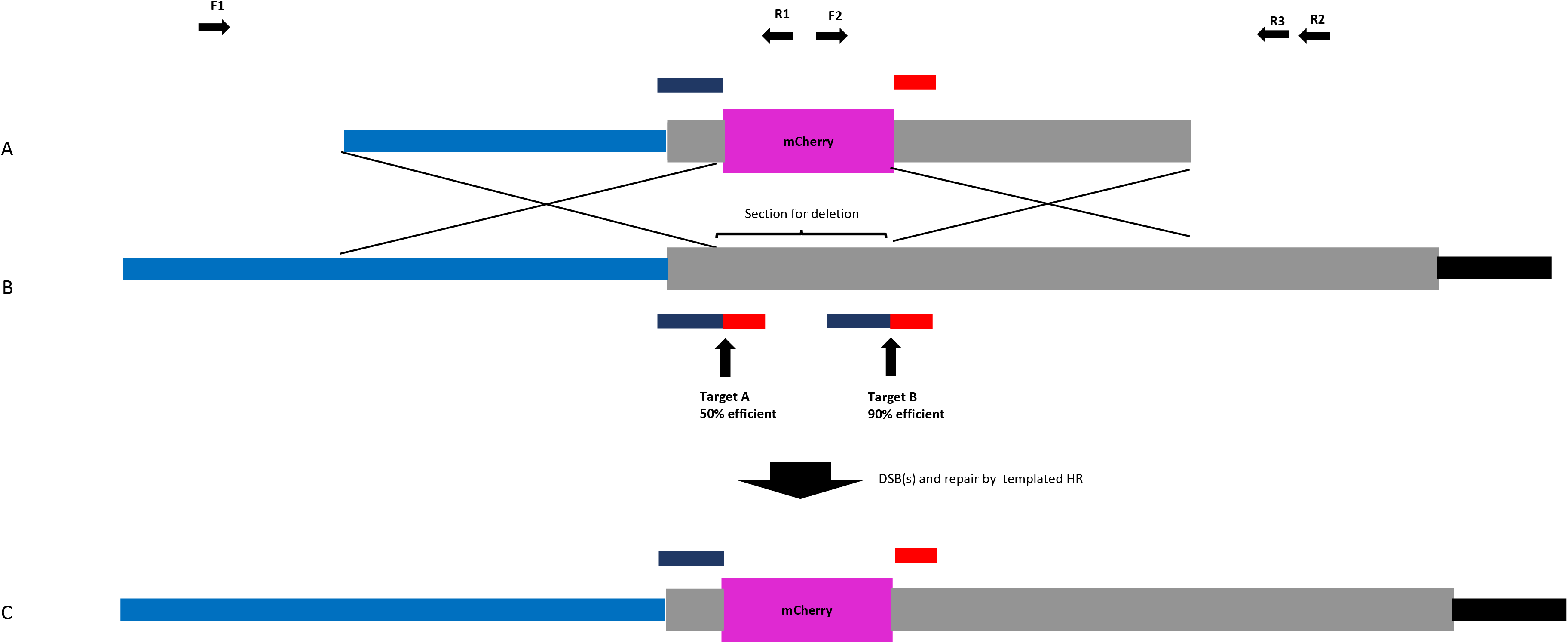
**A** repair template, **B** target locus, **C** successful GT into target locus. Promoter region of HORVU4Hr1G061310 in light blue, mCherry CDS in purple, HORVU4Hr1G061310 CDS in grey and 3’UTR in black. Protospacer sequences A and B in dark blue. Associated PAMs in red. Guide efficiency is shown as % of lines in which indels detected. Complete protospacer/PAM sequences are absent from A and C to prevent cleavage by Cas9. A successful event leads to a partial deletion of the HORVU4Hr1G06131 CDS with the remainder being fused in-frame to mCherry. Forward (F) and reverse (R) screening primers are indicated as black horizontal arrows.

**Figure 1.2.**
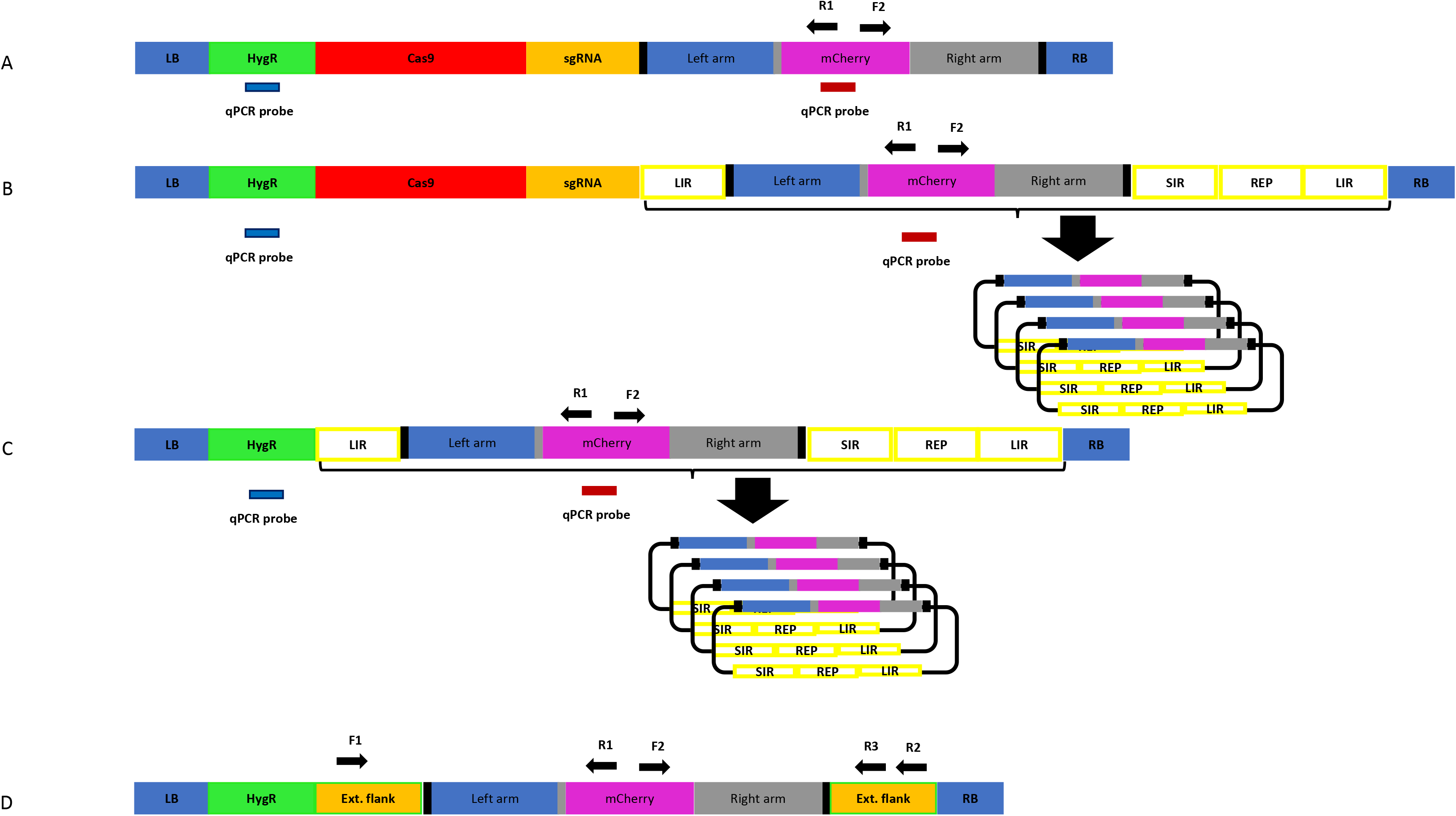
Constructs for plant transformation used in this study: **RB** right border, **LB** left border, **HygR** hygromycin resistance cassette,**Cas9** *Hs* codon optimised *Sp*Cas9 driven by ZmUbiquiutin promoter, **sgRNA** guide RNA expression cassette for two guides (A & B), **Left arm** left homology arm, **mCherry** mCherry reporter CDS, **Right arm** right homology arm, **LIR** long intergenic region, **SIR** short intergenic region, **REP** replicase proteins, **qPCR probe** used for copy number determination. Thick black vertical bars indicate target sites for the Cas9/guides. Construct **A** is the basic GT version, construct **B** is the same except the repair template is contained between replicon sequences. Construct C is the same as B but lacks the Cas9/gRNA and ability to introduce DSBs. Construct **D** has no Cas9/gRNA or replicon but has the repair template with extended homology arms. It was transformed into barley and used to establish a sensitive PCR assay. **F1, R1, F2, R2, R3** are PCR primers used for screening.

Wheat Dwarf virus (WDV) is one of the Geminivirus family, members of which have been used in both dicotyledonous and monocotyledonous plants previously, as replicons to deliver genome editing reagents and in particular the repair template for GT (Baltes et al., 2014; Butler et al., 2016; Gil-Humanes et al., 2017; Wang et al., 2017; Dahan-Meir et al., 2018; Hahn et al., 2018; Vu et al., 2020). The coat and movement protein coding sequence can be completely removed and replaced by a fragment of no maximally determined size whilst still retaining the ability to replicate within its host to high copy numbers after introduction by *Agrobacterium* or physical means. We used the basic template (LIR-SIR-REP-LIR) previously used with success (Gil-Humanes et al., 2017; Wang et al., 2017) but supplemented for our purposes by using the genome sequence from a strain of WDV isolated from barley (WDV-Bar [Hu]) (Ramsell et al., 2009). The WDV-Bar[Hu] version of LIR-SIR-REP-LIR was included in construct B (figure 1.2b) such that it would allow rolling circle replication of the repair template already present in construct A. Previously such replicons have often been shown functional in terms of replicative ability by using PCR to detect the circular replicating form of the linearly supplied unit. We chose to develop a qPCR copy number assay using amplicon/probe combinations in the repair template, hygromycin selection cassette (figure 1.2) and a single copy barley gene to enable quantification of replication in stable transgenic lines.

Construct C (figure 1.2c) was identical to B other than lacking the Cas9 and sgRNA cassettes, so able to amplify the repair template but unable to induce the site specific DSBs. This was to test the importance of targeted DSBs in GT which have been shown to be beneficial (Fauser et al., 2012) although not always essential (Terada et al., 2002).

### Design of assay for detection of GT events

Construct D (figure 1.2d) was produced as a means of optimising the PCR screening strategy for GT detection, as high sensitivity and specificity would be vital due to the rarity of GT events and the expectation that we could be searching for somatic sectors which might represent a small proportion of the cells within leaf samples taken for analysis (Schiml et al., 2017). Construct D contains the repair template as found in constructs A, B and C however the homology arms have been extended for a few hundred nucleotides with the native HORVU4Hr1G061310 genomic sequence to include the binding sites for the F1, R2 and R3 primers (figure 1.2d). By creating a single copy transgenic line with construct D, as determined by qPCR assay, a more realistic scenario to derive template for optimisation was possible than by using plasmid alone. In order to allow distinction from true GT events, polymorphisms at the junctions of the extended flanks and the homology arms were introduced which would not be present in the predicted true GT events. Various PCR conditions were tried and the best (see methods) were found to work well with primer combinations F1/R1, F2/R2 and F1/R3. The most sensitive were found to be junction PCRs F1/R1 and F2/R2 which would identify GT events at either the left or right junction respectively. By serially diluting 30ng of construct D genomic DNA, considering the 5.3Gbp haploid barley genome and the average weight of 650 Daltons per base pair it was possible to calculate the number of template copies in each PCR reaction and thus determine the threshold sensitivity. This was found to be in the region of 40 copies for the F1/R1 primer pair (Supplementary file 1), so theoretically capable of identifying a somatic sector containing the same number of cells with a GT event. PCR with primers F1/R3, although covering the entire GT event over both left and right junctions was less sensitive, presumably due to the greater amplicon size and the competitive tendency of the smaller WT allele to amplify and dominate the products (see figure2). The limit of detection for the F1/R3 amplicon was in the region of 1000 template copies (data not shown). For this reason, it was decided to use the more sensitive F1/R1 & F2/R2 junction combinations for screening primary transgenics where small somatic GT sectors were likely.

### Production and analysis of the T0 generation

Barley *cv* Golden Promise was transformed with constructs A, B, C and D using *Agrobacterium* delivery and selection of transgenic plants was done on hygromycin containing media (Hinchliffe and Harwood, 2019). Construct A yielded 14 T0 lines (1826 prefix) and construct B (2158 prefix) 39 T0 lines. These were screened and scored using the F1/R1 and F2/R2 primer pairs as well as being assayed by qPCR for their HptII (T-DNA) and mCherry (repair template) copy numbers. This data is given in file S2 which shows that in the case of construct A (1826) lines the copy number of HptII and mCherry correspond as expected for two single copy elements on a T-DNA. The 39 construct B (2158) lines however show an average of 7575 copies of mCherry whilst the HptII copy number remains largely one or two. This indicates that in many of the 2158 lines rolling circle replication is occurring giving rise to huge numbers of repair template copies. File S2 also shows the presence or absence of F1/R1 and F2/R2 PCR products of the correct size and for 1826 lines, 2/14 (14%) scored positive for both left and right junction PCRs whilst for 2158 lines, 22/39 (56%) scored positive for the same two PCRs.

To check the identity and fidelity of these PCR products, F1/R1 and F2/R2 products were purified and Sanger sequenced for the lines 2158-9-1, 2158-14-1, 1826-5-2, 1826-8-1 and found to be identical and as expected for perfect GT events (Supplementary file 3). As expected, construct C lines (2291 prefix) also generated many copies of repair template, but unexpectedly also produced correctly sized PCR products with primers F1/R1 and F2/R2 which is shown in supplementary file 2. In fact, 8/16 (50%) of the 2291 lines gave both left and right junction PCRs of the size indicative of a GT event and furthermore when purified and sequenced gave exactly the same sequence as seen with the 1826 and 2158 lines. Looking at the relation between mCherry copy number and the presence/absence of F1/R1 and F2/R2 PCR products (supplementary file 2) it was apparent that high numbers of repair template and PCR success were linked. Whilst this could mean that increasing the number of repair template copies was causing GT it could also indicate that false PCR positives were being triggered by the high number of repair templates produced by the replicon.

To test this latter idea plasmid DNA containing the repair template was mixed with wild type Golden Promise DNA (where GT could not have occurred) and F2/R2 PCR performed. Initially 30ng of barley DNA (as used in all other screening PCRs described) was mixed with around 7.72 × 109 copies of repair template and this resulted in the production of the 1047bp F2/R2 band. This plasmid was then titrated against the 30ng wild type barley DNA (representing 5240 target site copies) to determine the minimum number of repair template copies per target site necessary to trigger the false positive when 30ng of barley DNA was used as template. This is shown in file S4 and was found to be in the region of 700 copies per target site, based on the 5.3Gbp genome size and the average weight of a single base pair to be 650 Daltons. This result can be related to the qPCR copy number determinations for mCherry (repair template) in the replicon lines where the numbers in supplementary file 2 relate to copies per haploid genome or in other words per target site (there is one copy of HORVU4Hr1G061310 per haploid genome). Looking at supplementary file 2 it is evident that F1/R1 and F2/R2 products begin to appear in 2158 and 2291 lines at around 600 or 700 copies of mCherry per genome/target site, meaning it is likely that many of the PCR bands produced in replicon lines are false positives. This was further confirmed by sequencing a band from the plasmid titration test (file S4) in the lane labelled 736641 which proved identical in sequence to the F2/R2 bands obtained for the 2158, 2291 and 1826 lines. Presumably by increasing the number of repair template copies with the replicon we had inadvertently also increased the likelihood of partial primer extension from within the repair template. For example, R1 could in one cycle of PCR be partially extended from within mCherry to somewhere in the left homology arm. After denaturation, the partially extended product would be free to anneal at its 5’ end with the homologous site in the target region (template switching) where it could then be extended beyond the position of the F1 primer binding site. F1 could then prime against this site and extend to produce double stranded DNA of sequence identical to the predicted GT event and allow exponential amplification and production of the false positive.

The 1826 lines all had relatively low copy numbers of repair template (highest was 2), way below 600 per target site and so our testing indicated that the lines 1826-8-1 and 1826-5-2 would be true positives. Of course, the false positives in the replicon lines could be masking true positives in the background, so the 39 individual 2158 lines were subject to F1/R3 PCR along with the 14 individual 1826 lines, however none produced a band (data not shown) although this was unsurprising due to the low sensitivity of this large amplicon PCR. Accordingly lines 1826-8-1 and 1826-5-2 were sown out for T1 screening due to being likely true positive GT lines, whilst 17 F1/R1 & F2/R2 positive 2158 lines were selected for T1 screening based on the assumption that some true positive GT events may be masked by false positives created via replicon amplification.

### Analysis of the T1 generation and beyond

Because of the false positive PCR issue, and to detect GT events and somatic sectors of significant size likely to become heritable, it was decided to screen T1 plants with the less sensitive F1/R3 primer pair. For each of the 17 selected T0 2158 lines approximately 70 siblings were sown out, giving a total of around 1200 from which no F1/R3 positives were identified. T0 line 1826-5-2 produced 228 seeds and all were sown and screened producing no F1/R3 positive band. T0 line 1826-8-1 was however more productive and yielded 467 seeds and from these, 3 T1 plants produced a band of 2.2Kb indicative of the sought-after GT event as well as a second band of 1.6Kb corresponding to the wild type allele. These 3 T1 siblings were designated 1826-8-1_A, 1826-8-1_B and 1826-8-1_C. The 2.2Kb band was purified for all three siblings and sequenced from end to end, showing that all were identical to the predicted GT event (supplementary file 3). The three sibling T1 plants were grown to maturity and harvested before sowing out seeds for T2 screening. 94 individual 1826-8-1_A T2 siblings were screened and 75 gave the GT PCR product and 19 gave no band, corresponding to a 3:1 ratio (supplementary file 5), which is expected if the event was heterozygous in the T1 parent. Eight of these T2 siblings are shown in figure 2 after F1/R3 PCR where homozygous, heterozygous and wild type plants can all be clearly seen. This strongly indicates that the GT event occurred either in the T0 generation or very early in T1 i.e. just after fertilisation. All 94 of the 1826-8-1_A T2 siblings contained 2 copies of the T-DNA (homozygous) as determined by qPCR, so homozygous GT plants (figure 2) were selected for crossing to wildtype Golden Promise in order to segregate away the T-DNA from the GT event in F2.

**Figure 2.**
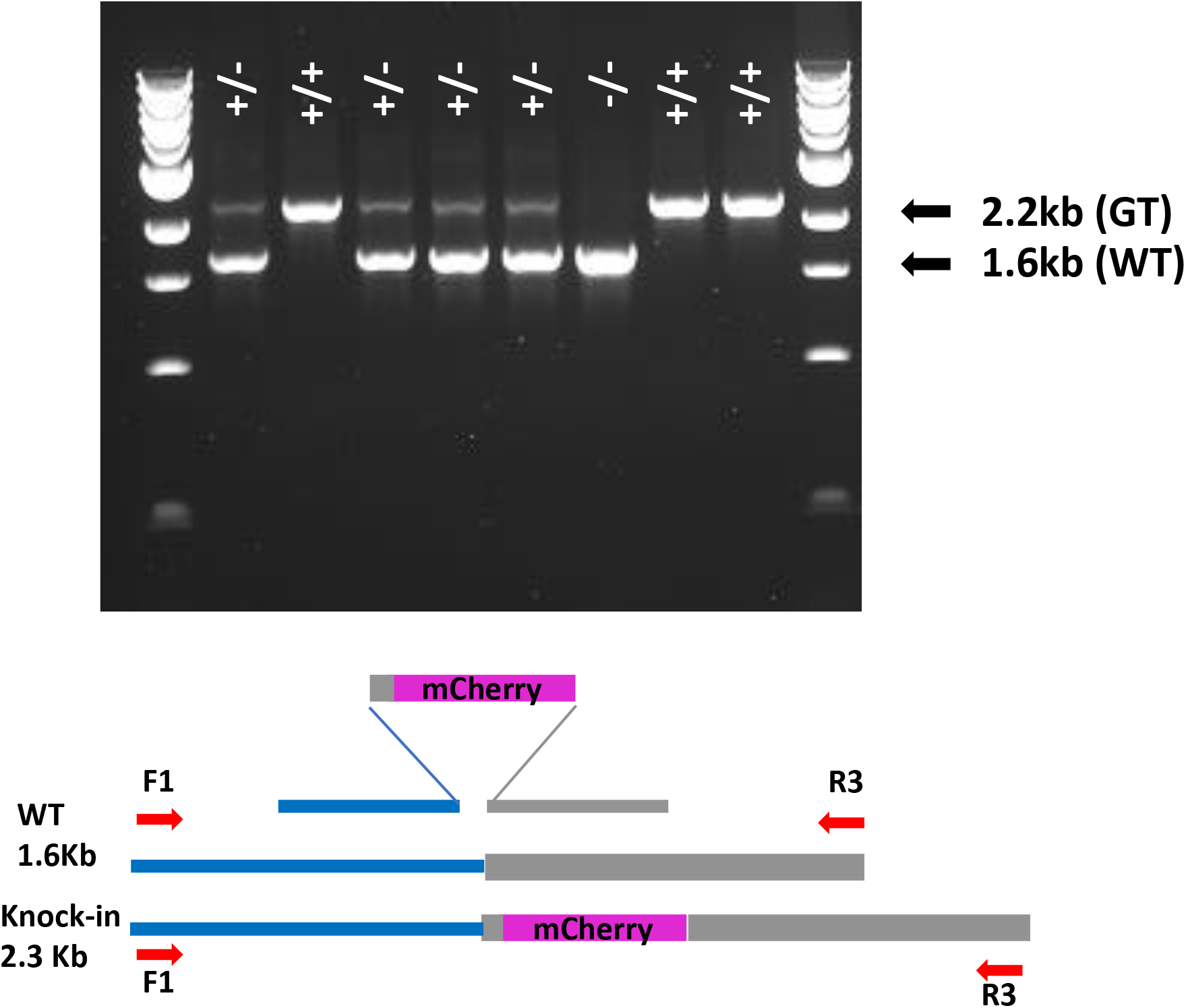
Gel showing segregation of GT event in 1826-8-1_A T2. Bands are the products of F1/R3 primers. + indicates GT allele, – indicates WT allele.

T1 line 1826-8-1_B was also sown out for T2 screening, but out of 94 siblings none screened positive for GT. This indicates that the event which was detected in T1 with F1/R3 primers and sequencing, would according to the sensitivity of the assay represent a somatic sector of at least 1000 cells, which was unable to pass through the germ line into T2 plants and was therefore lost. All 94 of the 1826-8-1_B T2 siblings contained 2 copies of the T-DNA (homozygous).

T1 line 1826-8-1_C was sown out for T2 screening and this time 11/94 screened positive for GT. All 94 of the 1826-8-1_C T2 siblings contained 2 copies of the T-DNA (homozygous). Three of the GT positive T2 siblings were designated 1826-8-1_C1, 1826-8-1_C2, 1826-8-1_C3 and sown out again for T3 screening. 17/24 1826-8-1_C1 siblings (3:1), 16/24 1826-8-1_C2 siblings (3:1) and 24/24 1826-8-1_C3 siblings were positive for GT indicating that T2 parents were likely GT heterozygotes (C1, C2) or GT homozygotes (C3) (see supplementary file 5). This is consistent with a T1 parent which was a cellular mosaic of the GT event(s) which passed through the germline into T2 progeny at a subsequently lower fraction than the 75% expected from a heterozygous parent. Alternatively, GT could have occurred independently in the T2 lines 1826-8-1_C1, 1826-8-1_C2 and 1826-8-1_C3 to give the same T3 GT zygosity.

As with line 1826-8-1_A, all T2 siblings of 1826-8-1_C were homozygous for the T-DNA insertion, so it was not possible to lose the transgene without backcrossing to Golden Promise. As all 1826-8-1 lines share the same T-DNA insertion and the crossing was already underway for 1826-8-1_A, this was not done for 1826-8-1_C.

### Linkage of the T-DNA and GT event

All 19 F1 lines produced for the 1826-8-1_A x Golden Promise cross were heterozygous for the T-DNA and GT as determined by qPCR and F1/R3 PCR. In F2 74/96 (3:1) siblings screened positive for GT as expected. All 96 were also tested for the presence of the transgene by qPCR which showed that all siblings containing the GT also contained the T-DNA and all GT free plants were also free of the T-DNA (supplementary file 5), in other words the T-DNA and GT locus are linked. To see how close the GT and T-DNA were to each other, a chromosome walking technique was used to determine the flanking sequences of the T-DNA. BLAST search using the sequence obtained as query against the barley genome revealed the T-DNA to be located 4.23Mb from the GT locus on chromosome 4 in line 1826-8-1 (supplementary file 6). The same chromosome walking was also done for line 1826-5-2 which was found to harbour the T-DNA on chromosome 7 (supplementary file 6).

## Discussion

Figure 3 summarises the key findings described above for all plants analysed. Heritable GT was confined to line 1826-8-1 with the event in 1826-8-1_A occurring either in T0 or very early T1 and the 1826-8-1_C events occurring in T1 or T2. Additionally, a significant event leading to detection with the low sensitivity primer pair F1/R3 was recovered in 1826-8-1_B but lost by T2 so must have occurred in T1. This shows that the 1826-8-1 family tree had diverged before the origin of these independent GT events and so for some reason the line 1826-8-1 was relatively prolific in terms of GT. A comparable line 1826-5-2 showed somatic GT in T0 but did not go on to result in subsequent heritable GT. This may be related to the T-DNA containing the repair template being linked to the target site in 1826-8-1 but not in 1826-5-2. It was previously reported that if the repair template and target site were present on the same chromosome then GT was around twice as frequent as when they were on different chromosomes (Fauser et al., 2012). Successful GT in line 1826-8-1 also makes sense in light of evidence that DNA repair by HDR using a sister chromatid template is common in barley (Vu et al., 2014). Being on the same chromosome is likely to impact on the physical proximity of target and donor site. It was recently reported in rice that using a Cas9-VirD2 fusion to direct the repair template to the target site had a beneficial effect on GT (Ali et al., 2020). It is also reported that the zygosity of the repair template has a similar impact (Puchta et al., 1995), where a homozygous transgene was 50% more likely to lead to intrachromosomal HR based gene repair than if hemizygous. In line with this, all three 1826-8-1 T1 siblings of interest were homozygous for the T-DNA whilst the overall T1 T-DNA inheritance in this line showed 3:1 segregation.

**Figure 3.**
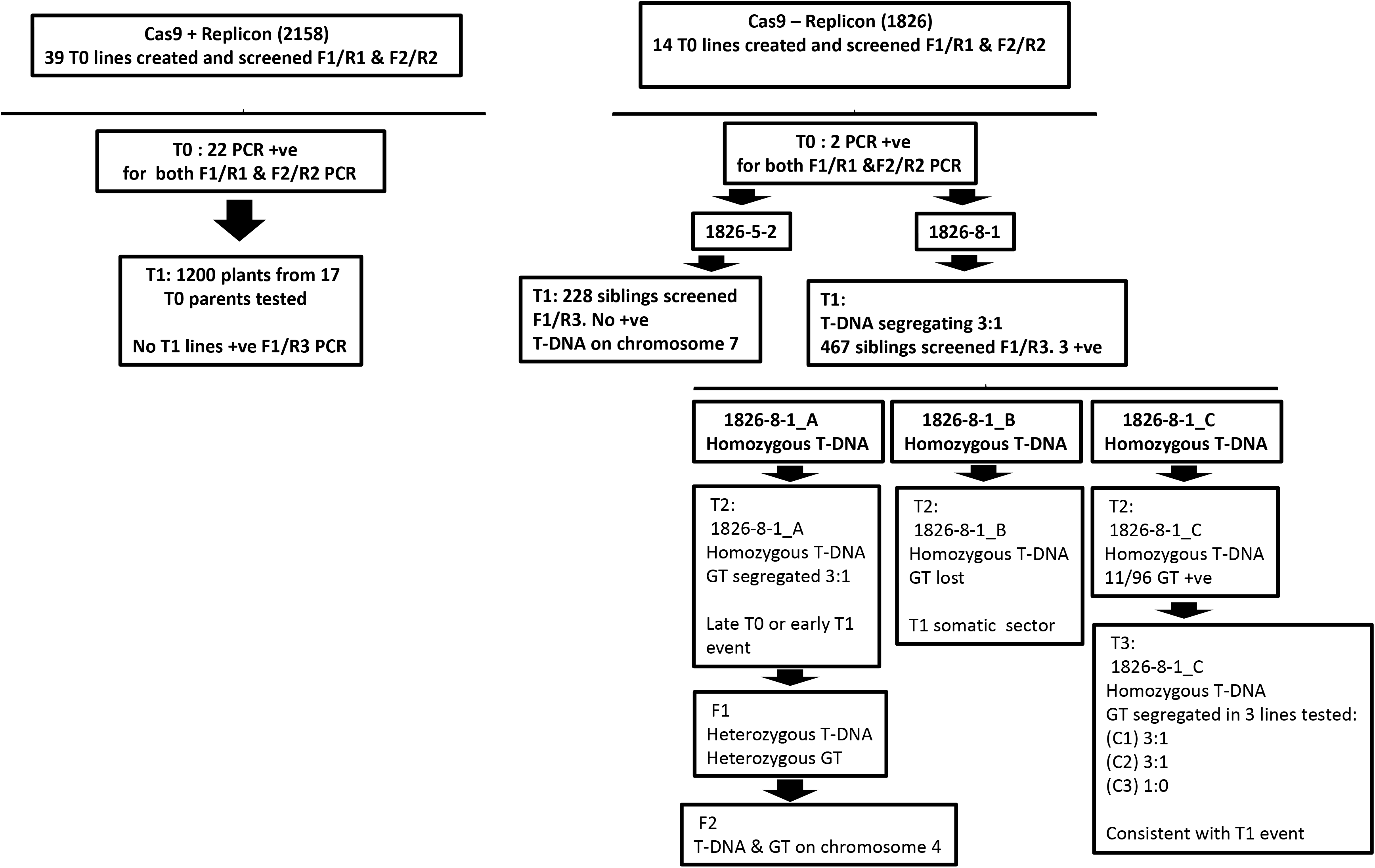
Key findings comparing in-planta GT with and without geminiviral replicon. 2158 are replicon lines (construct B) and 1826 lines are non-replicon lines (construct A).

We did not carry out any microscopy study to see if the in-frame mCherry HORVU4Hr1G061310 fusion created was functional as screening all 19 1826-8-1_A F1 plants produced showed that they still contained the T-DNA based repair template (data not shown). Similarly, 6 GT positive T3 plants from each of 1826-8-1_C1, 1826-8-1_C2 and 1826-8-1_C3 all contained the T-DNA based repair template (data not shown). This repair template contains the promoter region, mCherry and much of the HORVU4Hr1G061310 CDS so may well have given a fluorescent signal despite not being integrated at the target locus by GT. This would not allow distinction of a signal arising from GT at the target locus and a signal from the repair template still located in the T-DNA.

2158 lines had very high copy numbers of the repair template although none of the 17 T0 lines screened in T1 gave rise to significant or heritable F1/R3 detectable GT events despite screening twice the number of plants in T1 for the 2158 lines compared to 1826 lines. 2158 repair template amplification was detected in T0 and again in T1 (supplementary file 7) where replication co-segregated with the T-DNA. Despite there being no issue affecting heritability of a replicationally functional T-DNA, we have no data to show whether the repair template copy number was increased in the cells giving rise to the germ line. With this in mind, it could be possible that the replicon had a positive effect on GT in somatic cells not giving rise to the germ line. However titration of repair template plasmid against wild type Golden Promise DNA in vitro indicated that the GT activity detected in T0 2158 lines was potentially a PCR artefact as junction PCR bands begin to appear at around 700 copies of repair template per target site, which is very close to the ratio seen *in-planta* with the replicon where the junction PCR began yielding product. Future GT experiments utilising high copy numbers of repair template should be aware that such an approach is liable to produce false positive PCR results and would benefit from strategies to prevent them. One way to do this may be to reduce the length of homology arms to a minimum, thus reducing the size of the region in which partial primer extension may occur before template switching during PCR. Whilst reducing the length of homology arms may result in a decrease in overall GT efficiency, relatively short homology arms of 196bp and 74bp have been shown to function in rice (Li et al., 2020). Another way to reduce false positive junction PCR may be to simply increase the size of the amplicon by moving the primer in the flanking non-repair template region further out, which may in turn reduce the chances of a partially extended product being fully extended after template switching. From our repair template titration experiments it would clearly be a good idea to test the potential for false positive junction PCR by mixing WT genomic DNA and repair template in vitro before PCR.

It may be that in our experiments, GT was occurring in replicon lines in the T0 generation but was indistinguishable from the false positive junction PCRs being triggered and was perhaps at a low mosaic density, or not in pre-germinal tissue, and so insufficient to allow its inheritance into T1. Although we screened a greater number of T1 progeny (1200>695) from a greater number of T0 parents (17>2) for the replicon (2158) compared to the non-replicon (1826) lines, we cannot be sure that this is a valid replicon/non-replicon comparison as indel formation at target sites may have been unequal for some reason. Using Cas9 has the potential disadvantage that indel formation is likely to mutate the “seed region” of the target site such that further DSBs are not possible as the relevant guide no longer matches the site. We know that both guide A and B were able to induce indels in 50% and 90% respectively of T0 lines as detectable by PCR and direct Sanger sequencing, which is a relatively insensitive assay and may well indicate that many target sites would no longer be available for GT. As we only had one non-replicon (1826) T0 transgenic that yielded heritable GT events, a larger number of primary transgenics and GT events would need to be investigated in order to make a replicon/non-replicon comparison. Additionally, the target sites in T0,T1,T2, etc. lines created could be sequenced to gain more insight into the remaining availability of WT target sites.

Recently it has been shown in *Arabidopsis* that timing the occurrence of DSBs to the egg cell greatly increases GT efficiency (Miki et al., 2018; Wolter et al., 2018). Similarly, by using Cas12a instead of Cas9 GT efficiency was increased (Wolter and Puchta, 2019). Two features here address the potential lack in availability of WT target sites that may be shutting down DSB formation in our experiment. Firstly, restricting DSBs to egg cells would mean each female gamete has the potential for DSBs to occur and in turn undergo GT, rather than a reduced or non-existent fraction resulting from indels formed earlier during development under ubiquitous Cas9 expression. Secondly, Cas12a cuts outside of its seed region and would be expected to resist a certain amount of indel formation and may therefore keep creating DSBs for an increased length of time compared to Cas9, giving more potential for GT to occur. It will be interesting to see if the benefits to GT of egg cell specific Cas12a can be translated to crops.

A previous report of in-planta GT in *Arabidopsis* (Hahn et al., 2018) found no beneficial effect from including the repair template within a replicon, whilst a single copy repair template (similar to our construct A) gave rise to inheritable GT. However, this study investigated the progeny of just three primary transformant lines per DNA construct and may also suffer from indels shutting down target sites. In tomato, bean yellow dwarf virus-based replicons have been shown to result in heritable GT events (Cermak et al., 2015; Dahan-Meir et al., 2018; Vu et al., 2020). In one tomato study utilising an in-planta approach, it increased the percentage of inheritable T0 events from 8% without a replicon to 25% with a replicon (Dahan-Meir et al., 2018). Rice (Wang et al., 2017), wheat (Gil-Humanes et al., 2017) and potato (Butler et al., 2016) replicon/GT reports describe junction PCR/sequencing assays similar to our false positive prone F1/R1 and F2/R2 and no GT heritability.

Our work in barley has extended what has previously been shown in this species as we created the first heritable true GT events at a native locus. However, we were unable to segregate away the editing reagents on the T-DNA, possibly due to an inadvertent selection for linkage. Whilst it may be possible to separate the two loci by searching for meiotic recombinants this probably represents an unreasonable amount of work. Increasing the number of heritable GT events detected will probably allow the isolation of unlinked versions which would in turn be easier if GT efficiency was boosted in other ways, such as egg cell Cas12a expression. Additionally, a pooling strategy may enable more plants to be screened which should increase the numbers of GT events recovered.

## Conflict of Interest

The authors declare that the research was conducted in the absence of any commercial or financial relationships that could be construed as a potential conflict of interest.

## Author Contributions

TL designed the experiments, conducted molecular work and analysis and prepared the manuscript. AH carried out barley transformations. MC carried out the crossing programme. YM assisted with molecular analysis and crossing. WH contributed to the study design and manuscript preparation. All authors read and approved the final manuscript.

## Acknowledgments

We thank JIC horticultural services for plant care. We acknowledge support from the project Engineering Nitrogen Symbiosis for Africa (ENSA) currently supported through a grant to the University of Cambridge by the Bill & Melinda Gates Foundation and the Foreign, Commonwealth and Development Office (FCDO). The work was also supported by the UK Biotechnology and Biological Sciences Research Council (BBSRC) through grant BB/P013511/1.

## Supplementary Material

### List of supplementary files

#### Supplementary file 1

Gel showing sensitivity obtained in F1/R1 PCR screen setup. Construct D was transformed into barley and DNA extracted from a regenerated plant (transgene copy number 1) and quantified by Qubit fluorescence. Serial dilutions were made of this DNA for subsequent PCR. The copy number of transgene D are shown for each lane. The limit of detection is around 40 copies of the target.

#### Supplementary file 2

qPCR determined copy numbers of HptII (TDNA), mCherry (repair template) and presence/absence of junction PCR products (right:F1/R1; left: F2/R2) for T0 lines.

#### Supplementary file 3

Sequences of GT events for lines 2158-9-1, 2158-14-1, 1826-5-2, and 1826-8-1 showing F1/R1 (T0), F2/R2 (T0) and F1/R3 (T1) products.

#### Supplementary file 4

Copies of repair template plasmid per target site. 30ng of wild type barley DNA was mixed with serial dilutions of plasmid DNA containing the repair template. The lane numbers represent the copies of repair template: target site ratio. Positive control was a construct D line. The 1047bp band in lane 736641 was excised, purified and sequenced.

#### Supplementary file 5

Table showing zygosity of T-DNA and GT events in interesting descendants of 1826-8-1 throughout the generations analysed in this study.

#### Supplementary file 6

Flanking sequences of the T-DNA in lines 1826-8-1 and 1826-5-2.

#### Supplementary file 7

T1 inheritance of the T-DNA and associated replicon activity.

#### Supplementary file 8

Primer and probe sequences used in the study and sequences of construct components.

## Notes

### Competing Interest Statement

The authors have declared no competing interest.

