## Supplementary files for "In-planta Gene Targeting in Barley using Cas9, with and without Geminiviral Replicons"

### File S1

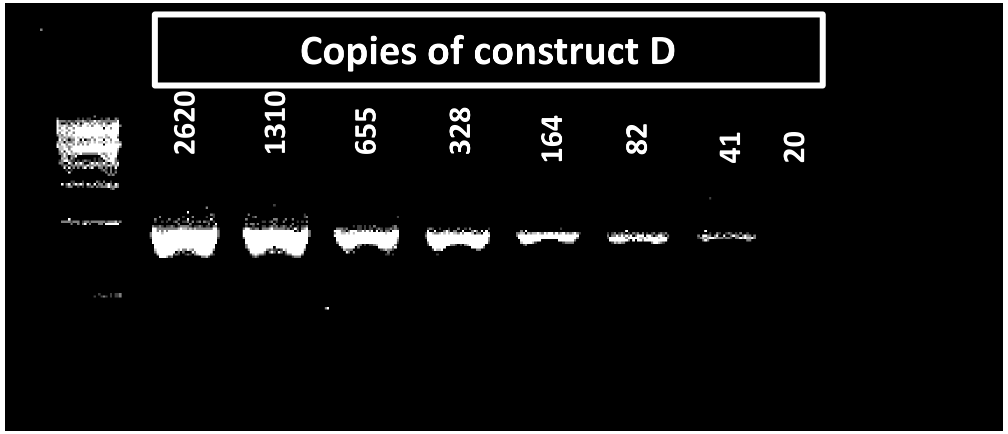

**File S1**

Gel showing sensitivity obtained in F1/R1 PCR screen setup. Construct D was transformed into barley and DNA extracted from a regenerated plant (transgene copy number 1) and quantified by Qubit fluorescence. Serial dilutions were made of this DNA for subsequent PCR. The copy number of transgene D are shown for each lane. The limit of detection is around 40 copies of the target.

**Table S2 showing qPCR determined copy numbers of HptII (TDNA), mCherry (repair template) and presence/absence of junction PCR products (right:F1/R1; left: F2/R2) for T0 lines**

| Line | HptII copies/diploid genome | mcherry copies/diploid genome | Right PCR | Left PCR |
|---|---|---|---|---|
| 1826-3-1 | 1 | 1 | NO | YES |
| 1826-5-2 | 2 | 2 | YES | YES |
| 1826-2-2 | 1 | 1 | NO | NO |
| 1826-2-1 | 1 | 1 | NO | NO |
| 1826-1-1 | 2 | 2 | NO | NO |
| 1826-6-1 | 1 | 1 | NO | NO |
| 1826-5-1 | ND | ND | NO | NO |
| 1826-4-1 | 1 | 1 | NO | YES |
| 1826-8-2 | 1 | 1 | NO | NO |
| 1826-10-1 | 1 | 1 | NO | NO |
| 1826-8.1 | 1 | 1 | YES | YES |
| 1826-9-1 | 1 | 1 | NO | NO |
| 1826-7-1 | 1 | 1 | NO | NO |
| 1826-6-2 | 1 | 1 | NO | NO |
| **Line** | **HptII copies/diploid genome** | **mcherry copies/haploid genome** | **Right PCR** | **Left PCR** |
| 2158_5_1 | 1 | 20 | NO | NO |
| 2158_6_1 | 1 | 5563 | YES | YES |
| 2158_7_1 | 1 | 38395 | YES | YES |
| 2158_8_1 | 1 | 436 | NO | NO |
| 2158_10_1 | 1 | 4 | NO | NO |
| 2158_9_1 | 16 | 85314 | YES | YES |
| 2158_4_2 | 1 | 888 | NO | YES |
| 2158_1_2 | 2 | 15 | NO | NO |
| 2158_2_2 | 1 | 3668 | YES | YES |
| 2158_3_2 | 3 | 7 | NO | NO |
| 2158_3_3 | 3 | 559 | NO | NO |
| 2158_11_1 | 1 | 3363 | YES | YES |
| 2158_1_1 | 2 | 4407 | YES | YES |
| 2158_2_1 | 1 | 784 | YES | YES |
| 2158_3_1 | 4 | 4008 | YES | YES |
| 2158_4_1 | 1 | 673 | YES | YES |
| 2158_12_1 | 2 | 4065 | YES | YES |
| 2158_18_2 | 1 | 5 | NO | NO |
| 2158_14_1 | 2 | 6327 | YES | YES |
| 2158_19_1 | 2 | 5494 | YES | YES |
| 2158_15_2 | 1 | 5028 | YES | YES |

| | | | | |
|---|---|---|---|---|
| 2158_13_1 | 100 | 608 | YES | NO |
| 2158_14_2 | 1 | 1276 | YES | NO |
| 2158_20_2 | 1 | 6127 | YES | YES |
| 2158_20_1 | 1 | 1264 | YES | YES |
| 2158_16_1 | 1 | 6048 | YES | YES |
| 2158_21_1 | 1 | 1 | NO | NO |
| 2158_22_3 | 1 | 287 | NO | NO |
| 2158_18_1 | 1 | 26375 | YES | YES |
| 2158_12_5 | 2 | 26076 | YES | YES |
| 2158_21_4 | 2 | 3037 | YES | YES |
| 2158_22_1 | 6 | 12957 | YES | NO |
| 2158_17_1 | 4 | 8665 | YES | YES |
| 2158_15_4 | 4 | 28568 | YES | NO |
| 2158_9_3 | 1 | 1 | NO | NO |
| 2158_23_2 | 1 | 1370 | YES | YES |
| 2158_24_2 | 1 | 2340 | YES | NO |
| 2158_7_3 | 1 | 1420 | YES | YES |
| 2158_17_4 | 1 | 4 | NO | NO |
| 2291-5-2 | 2 | 9489 | YES | YES |
| 2291-5-1 | 1 | 8 | NO | NO |
| 2291-4-1 | 1 | 181 | NO | NO |
| 2291-3-1 | 1 | 33744 | YES | YES |
| 2291-2-1 | 3 | 602 | NO | NO |
| 2291-1-1 | 1 | 377 | NO | NO |
| 2291-16-1 | 1 | 990 | YES | NO |
| 2291-15-1 | 2 | 2 | NO | NO |
| 2291-14-1 | 5 | 36582 | YES | YES |
| 2291-13-1 | 3 | 52203 | YES | YES |
| 2291-12-1 | 6 | 4003 | YES | YES |
| 2291-11-1 | 2 | 32024 | YES | YES |
| 2291-10-1 | 500 | 5726 | YES | YES |
| 2291-9-1 | 2 | 2 | NO | NO |
| 2291-8-1 | 300 | 10371 | YES | YES |
| 2291-7-2 | 1 | 220 | NO | NO |
| 2291-7-1 | 1 | 397 | NO | NO |

ND = Not determined

**Supplementary file S3**

Sequences of GT events for lines 2158-9-1, 2158-14-1, 1826-5-2, 1826-8-1

F1/R1 T0 products

agaggttagcctttgtagatggtattgggcttatattgggttccatgggcggagtttgtttttcgaacaccagatttataggcatggtaaatcaaat
gtttttatggcacatatgtgttcgctgaggatggcaagtttagttaacaagcatgtcaaatttgactcaatttatttttgccaagaaattgtcgtgct
tgcaaacttaatttgccaccctgacgccaatataaattgccatacaaaatgtttgatttgccatgcctaattttttagcatttttgtttcttttatcatt
agccattatcttcttctttttaaaatcttatagtatgcaaatctaataagattcctggcgagttattaagaaaaccagattctcattatttttttcctttg
caaaaaagagaagattctctctctctcttacaacgattctcattctccggctcaaaaaaaagtttgtttctcattctcttcctgcttaatgcaatcggt
atttttttttgaggggaacttaatgcaatcagtagagtgcttgcctcgttgctggaaaaagaattgatgatccatgtgatttaacgagaaaaaca
aagtccgcccatggtgcccaatattttaggcccagttgggatggtagaacctgctgctggagccagctccagttttgttcgccgaatgccgagtcc
cggcgcccagggatggctataaataagcgagctcccgtgtccttgtgtacttgtaaaatctgtgctccctgcccaccgctctcccctcggttcccac
gcgccaaaacattccaacgtggagacaagaagcagcatagcgtgacaacgagggagggagccatggacgtgaccatggaggacgtgatggt
gagcaagggcgaggaggataacatggccatc

F2/R2 T0 products

gaccacctacaaggccaagaagcccgtgcagctgcccggcgcctacaacgtcaacatcaagttggacatcacctcccacaacgaggactacac
catcgtggaacagtacgaacgcgccgagggccgccactccaccggcggcatggacgagctgtacaagctccacggcctgatcgagtccatgct
ctgcgatgacactctcatcggcacgcccgagcccgacgagcacccagacccagccatgttcacggacggcccctgctactccaacggctccgac
ccgagcagcaccaccacgacgaacccgggcacgcccgtgcagcacgacgacgacctgccgcaggactgcaatcccgagaagggactccggct
gcttcacctgctcatggccgccgccgaggcgctctccggcccgcacaagagccgggagctggcacgggtgatattggttcggctcaaggagatg
gtctccagcaccagcggcaacgctgccgcgtccaacatggagcgcctcgccgcccacttcaccgacgcgctccaggggctcctcgatgggtccc
actccgtcgctgggaccagcaggcaggccgcatcccaccaccacagcaccggcgacgtgttgacggcattccagatgctccaggacatgtcgcc
ctacatgaagttcggccacttcaccgcgaaccaggcgatcctggaggcggtggcgggcgaccggcgcgtccacatcgtggactacgacctcgcc
gagggcatccagtgggcgtccctgatgcaggctatgacatcacgacccgatggcgtgtcgcctccgcacctgcgtatcaccgccatcacgcgga
gtggcgggggcggcgcgcgggcagtccaggaggccggacggcgcctcgcggccttcgcggggtccatcgggcagcccttctcgttcggacattg
ccgtctggactcggacgagaggttccggccggcgaccgtcaggatggtcaagggggagacgctcgtggcca

F1/R3 T1 products

agaggttagcctttgtagatggtattgggcttatattgggttccatgggcggagtttgtttttcgaacaccagatttataggcatggtaaatcaaat
gtttttatggcacatatgtgttcgctgaggatggcaagtttagttaacaagcatgtcaaatttgactcaatttatttttgccaagaaattgtcgtgct
tgcaaacttaatttgccaccctgacgccaatataaattgccatacaaaatgtttgatttgccatgcctaattttttagcatttttgtttcttttatcatt
agccattatcttcttctttttaaaatcttatagtatgcaaatctaataagattcctggcgagttattaagaaaaccagattctcattatttttttcctttg
caaaaaagagaagattctctctctctcttacaacgattctcattctccggctcaaaaaaaagtttgtttctcattctcttcctgcttaatgcaatcggt
atttttttttgaggggaacttaatgcaatcagtagagtgcttgcctcgttgctggaaaaagaattgatgatccatgtgatttaacgagaaaaaca
aagtccgcccatggtgcccaatattttaggcccagttgggatggtagaacctgctgctggagccagctccagttttgttcgccgaatgccgagtcc
cggcgcccagggatggctataaataagcgagctcccgtgtccttgtgtacttgtaaaatctgtgctccctgcccaccgctctcccctcggttcccac
gcgccaaaacattccaacgtggagacaagaagcagcatagcgtgacaacgagggagggagccatggacgtgaccatggaggacgtgatggt
gagcaagggcgaggaggataacatggccatcatcaaggagttcatgcgcttcaaggtgcacatggagggctccgtgaacggccacgagttcga
gatcgagggcgagggcgagggccgcccctacgagggcacccagaccgccaagctgaaggtgaccaagggtggcccccctgcccttcgcctggg
acatcctgtcccctcagttcatgtacggctccaaggcctacgtgaagcaccccgccgacatccccgactacttgaagctgtccttccccgagggct
tcaagtgggagcgcgtgatgaacttcgaggacggcggcgtggtgaccgtgacccaggactcctccctgcaggacggcgagttcatctacaaggt
gaagctgcgcggcaccaacttcccctccgacggcccagtaatgcagaagaaaaccatgggctgggaggcctcctccgagcggatgtaccccga
ggacggcgccctgaagggcgagatcaagcagaggctgaagctgaaggacggcggccactacgacgctgaggtcaagaccacctacaaggcc
aagaagcccgtgcagctgcccggcgcctacaacgtcaacatcaagttggacatcacctcccacaacgaggactacaccatcgtggaacagtac
gaacgcgccgagggccgccactccaccggcggcatggacgagctgtacaagctccacggcctgatcgagtccatgctctgcgatgacactctca
tcggcacgcccgagcccgacgagcacccagacccagccatgttcacggacggcccctgctactccaacggctccgacccgagcagcaccacca
cgacgaacccgggcacgcccgtgcagcacgacgacgacctgccgcaggactgcaatcccgagaagggactccggctgcttcacctgctcatgg

ccgccgccgaggcgctctccggcccgcacaagagccgggagctggcacgggtgatattggttcggctcaaggagatggtctccagcaccagcg
gcaacgctgccgcgtccaacatggagcgcctcgccgcccacttcaccgacgcgctccaggggctcctcgatgggtcccactccgtcgctgggacc
agcaggcaggccgcatcccaccaccacagcaccggcgacgtgttgacggcattccagatgctccaggacatgtcgccctacatgaagttcggcc
acttcaccgcgaaccaggcgatcctggaggcggtggcgggcgaccggcgcgtccacatcgtggactacgacctcgccgagggcatccagtggg
cgtccctg

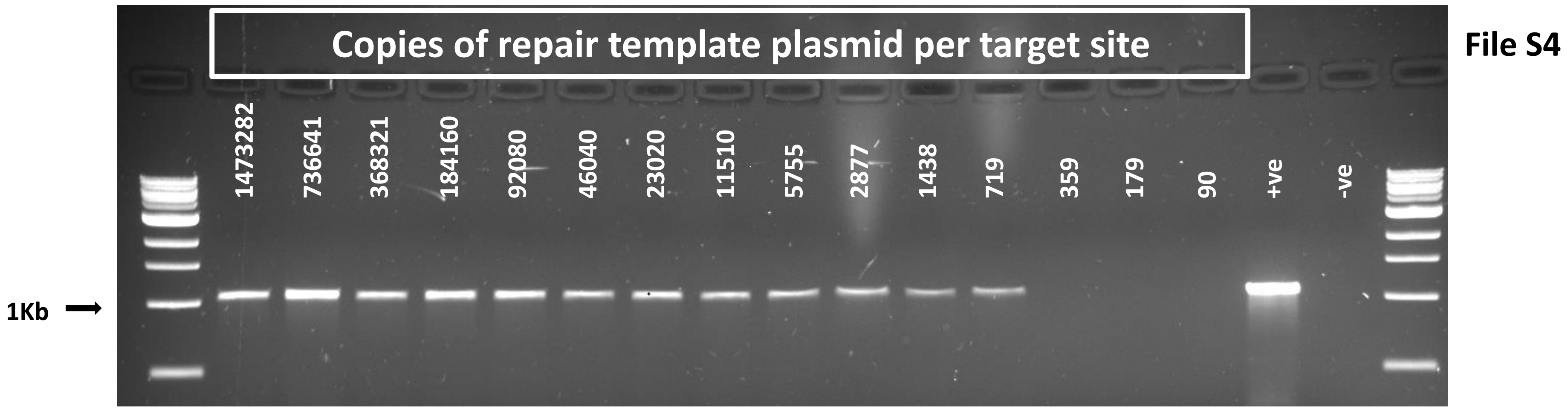

**File S4**
30ng of wild type barley DNA was mixed with serial dilutions of plasmid DNA containing the repair template. The lane numbers represent the copies of repair template: target site ratio. Positive control was a construct D line. The 1047bp band in lane 736641 was excised, purified and sequenced.

**Table showing zygosity of T-DNA and GT events in interesting descendants of 1826-8-1 throughout the generations analysed in this study**
T-DNA analysis by qPCR. GT analysis by PCR
* T-DNA and GT co-segregated

Segregation is confirmed by Chi square

| | T-DNA copies | | | | | | GT | | | | | |
|---|---|---|---|---|---|---|---|---|---|---|---|---|
| | T0 | T1 | T2 | T3 | F1 | F2 | T0 | T1 | T2 | T3 | F1 | F2 |
| **Sib. A** | 1 | 2 | 2 | | 1 | 3:1*(74/22) | Detected | Detected | 75/94 (3:1) | | Heterozygous | 3:1*(74/22) |
| **Sib.B** | 1 | 2 | 2 | | | | Detected | Detected | 0/94 (Lost) | | | |
| **Sib.C** | 1 | 2 | 2 | 2 | | | Detected | Detected | 11/94 | | | |
| **1826-8-1_C1** | | | | | | | | | | 17/24 (3:1) | | |
| **1826-8-1_C2** | | | | | | | | | | 16/24 (3:1) | | |
| **1826-8-1_C3** | | | | | | | | | | 24/24 | | |

| | Chromosome position | Flanking sequence obtained |
|---|---|---|
| **Target position** | HORVU4Hr1G061310/Ch 4 515362051 | |
| **1826-8-1 T-DNA** | Ch4 491249061. 752bp | ACTTGTCGCCTCGATCTCCGCCTTGATAGCCTCGATTTCCGTTCCTTCCCTTTAAGCAATGAGA<br>GAAGAAATGGGGGGGGGGGAAATCCCGTCACCTACCTCCATCTCTTCCTCGTCTTCCTCGCG<br>CTCTTCCTCTTCGAGCACCAGCGCCTCGACGTCGTCACCGACTCCAGGAGTTCCTCCATCGAC<br>CGTGGTCTTCCGGGCGCCAAGAAGCACGAAGTACTGCGCTTTCGTCCTCGGCACCGATCCTCT<br>TTTCCCAGGTACACACCTTCCCCCATTAGGGTTAGGCCGAGAACACCATGACCATTACTTCCT<br>GCGAGCTAGCCTTCATGACGCGACAAACACCCTACTTTAGATCCATTTCATATTTTTGGAATA<br>GACTCTTAGGCAGTAGCAAACTACGACCACTTTATCACATCTTTTCGTTAGCTTAGACCTTCAT<br>GCCGTAGATCCAATGTTGTCGCTGCGTTATGTGCATAACTCGCTACCATTATTTCAAGCAATAT<br>CCTTCTATTTTGATCCACTCGTAGATCCATAGTTTACTGCCTGTTTTTAGAAACATGTTTGCTCG<br>TCTCTCGAGAATATCCCTAAGGAAAAAGAACCCCTGCGAGTCGCCGTTTATCTCCAGCTTCCA<br>TAGCGACTTAAATTTATCACCATAGCCGTGACCCTTCTCTTTCCAATTTGAAGGGCGAGTTACC<br>ATACTTCAAAGATACTTCCATGAATCGCCCGAATGGCGGGTTATGATGTATGTCACCCATTTG<br>GCGGGTCACTAAATCCTTTTCGAACATACCGTAGACCCATACCGTCGAGTTAGAATACTTTCT |
| **1826-5-2 T-DNA** | Ch7 553846953. 310bp | AAAGCCCTAAACCTCTAGTATATAAAGGAGGAGGGAGGGGCAGGCCTTGCGCCCAAAGGGA<br>GCCTCCCTTGGGTCGGCCAAAGTGGAGGAGGGAGGAGTCCTCCTCCAACTCCACTTGGATTA<br>GGACTCTTTCCTTAATTTCCCACCTCTTTTGGATTTTCCACTTTTTCCTCATGGGCTTTCCTTGG<br>ATGACTTTGTCAGCCCATTATGCCTTATTGCACATCCAATAAACCCATGTGGACCCCTTGGGG<br>CGTGGTGGGCCCACCCAGTGGGCCCCCGGAACCCATTCGTCACTCCAGGTACACTGTCA |

**Table S7 showing T1 inheritance of T-DNA and associated replicon activity**

| Parent | HptII copies/diploid genome | mcherry copies/haploid genome |
|---|---|---|
| 2158_7_1 | 1 | 1104 |
| 2158_7_1 | 1 | 14669 |
| 2158_7_1 | 2 | 128780 |
| 2158_7_1 | 1 | 74142 |
| 2158_7_1 | 1 | 12717 |
| 2158_7_1 | 0 | 0 |
| 2158_7_1 | 0 | 0 |
| 2158_7_1 | 1 | 13274 |
| 2158_7_1 | 1 | 1396964 |
| 2158_7_1 | 1 | 1923 |
| 2158_7_1 | 2 | 32879 |
| 2158_7_1 | 0 | 0 |
| 2158_7_1 | 1 | 2753 |
| 2158_7_1 | 2 | 27301 |
| 2158_7_1 | 2 | 359261 |
| 2158_7_1 | 0 | 0 |
| 2158_7_1 | 1 | 6101 |
| 2158_7_1 | 1 | 39879 |
| 2158_7_1 | 1 | 101830 |
| 2158_7_1 | 1 | 8390 |
| 2158_7_1 | 0 | 0 |
| 2158_7_1 | 1 | 9676 |
| 2158_7_1 | 0 | 0 |
| 2158_7_1 | 1 | 194099 |
| 2291-13-1 | 5 | 137 |
| 2291-13-1 | 0 | 0 |
| 2291-13-1 | 5 | 173 |
| 2291-13-1 | 5 | 29 |
| 2291-13-1 | 0 | 0 |
| 2291-13-1 | 4 | 31311 |
| 2291-13-1 | 3 | 8490 |
| 2291-13-1 | 5 | 79 |
| 2291-13-1 | 3 | 613 |
| 2291-13-1 | 3 | 19774 |
| 2291-13-1 | 3 | 22991 |
| 2291-13-1 | 3 | 3186 |
| 2291-13-1 | 1 | 9 |
| 2291-13-1 | 1 | 31 |
| 2291-13-1 | 1 | 22 |
| 2291-13-1 | 3 | 2 |

| | | |
|---|---|---|
| 2291-13-1 | 1 | 11 |
| 2291-13-1 | 1 | 11 |
| 2291-13-1 | 1 | 2 |
| 2291-13-1 | 1 | 11 |
| 2291-13-1 | 3 | 12 |
| 2291-13-1 | 1 | 6 |
| 2291-13-1 | 0 | 0 |
| 2291-13-1 | 3 | 9 |

Supplementary file S8

| Copy number determination | | | |
|---|---|---|---|
| **Target** | **Forward primer** | **Reverse primer** | **Probe** |
| mCherry | GAGGCTGAAGCTGAAGGAC | GATGGTGTAGTCCTCGTTGTG | FAM-CCAACTTGATGTTGACGTTGTAGGCG-TAMERA |
| HptII | GGATTTCGGCTCCAACAATG | TATTGGGAATCCCCGAACATC | FAM-CAGCGGTCATTGACTGGAGCGAGG-TAMERA |
| Constans like | TGCTAACCGTGTGGCATCAC | GGTACATAGTGCTGCTGCATCTG | VIC-CATGAGCGTGTGCGTGTCTGCG-TAMERA |

| PCR primers | |
|---|---|
| F1 | TGCCAAAGTGCTACATCAGC |
| R1 | AAGCGCATGAACTCCTTGAT |
| F2 | CACTACGACGCTGAGGTCAA |
| R2 | GGCTTGGTGGAGTATGCAGT |
| R3 | CGTGATGTCATAGCCTGCAT |
| F4 | CAACGTGGAGACAAGAAGCA |
| R5 | TCCTTGAGCCGAACCAATATCAC |

| Sequencing primers | |
|---|---|
| Seq1 | TCAGCAGAGGTTAGCCTTTGTAG |
| Seq2 | CCTGGCGAGTTATTAAGAAAACCA |
| Seq3 | AGCCAGCTCCAGTTTTGTTC |
| Seq4 | CATCAAGGAGTTCATGCGCTT |
| Seq5 | GACGGCGAGTTCATCTACAAG |
| Seq6 | CAAGACCACCTACAAGGCCAA |

| | |
|---|---|
| Seq7 | ACCCAGACCCAGCCATGTT |
| Seq8 | CATCCCACCACCACAGCA |
| Seq9 | AGCCCTTCTCGTTCGGACATT |
| Seq10 | TGGTTTTCTTAATAACTCGCCAGG |
| Seq11 | AACATGGCTGGGTCTGGGT |

| **Chromosome walking** | | | |
|---|---|---|---|
| **Right border** | | **Left border** | |
| Rb_SP1 | TGAGACCGAGGATGCACATGTG | Lb_SP1 | ATTTCTTGTGTGCAACTCCGGGAA |
| Rb_SP2 | CATGTGACCGAGGGACACGAAGT | Lb_SP2 | GCCGTTTGTTGCCGCCTTTGTACAACCCCAGT |
| Rb_SP3a | AAGTGATCCGTTTAAACTNNNNNNNNNNNNCAGGAT | Lb_SP3a | CCCAGTCATCGTATATACNNNNNNNNNNNNCGTTAT |
| Rb_SP3b | ACGAAGTGATCCGTTTAANNNNNNNNNNNNTGACAG | Lb_SP3b | TATATACCGGCATGTGGANNNNNNNNNNNNACGTAG |

| **Construct assembly** | |
|---|---|
| Protospacer A | gtgaccatggaggacgtggt |
| Protospacer B | gacggcggccacgacctcca |

| | |
|---|---|
| Repair template | gacggcggccacgacctccacggaagcatgtcaaatttgactcaatttattttttgccaagaaattgtcgtgcttgcaaacttaatttgccaccctgacgccaatataaattgccatacaaaatgtttgat<br>ttgccatgcctaattttttagcatttttgtttcttttatcattagccattatcttcttctttttaaaatcttatagtatgcaaatctaataagattcctggcgagttattaagaaaaccagattctcattattttttc<br>ctttgcaaaaaagagaagattctctctctcttacaacgattctcattctccggctcaaaaaaaagtttgtttctcattctcttcctgcttaatgcaatcggtattttttttttgaggggaacttaatgcaatc<br>agtagagtgcttgcctcgttgctggaaaaagaattgatgatccatgtgatttaacgagaaaaacaaagtccgcccatggtgcccaatattttaggcccagttgggatggtagaacctgctgctggagcc<br>agctccagttttgttcgccgaatgccgagtcccggcgcccagggatggctataaataagcgagctcccgtgtccttgtgtacttgtaaaatctgtgctccctgcccaccgctctcccctcggttcccacgc<br>gccaaaacattccaacgtggagacaagaagcagcatagcgtgacaacgagggagggagccatggacgtgaccatggaggacgtgatggtgagcaagggcgaggaggataacatggccatcatca<br>aggagttcatgcgcttcaaggtgcacatggagggctccgtgaacggccacgagttcgagatcgagggcgagggcgagggccgcccctacgagggcacccagaccgccaagctgaaggtgaccaag<br>ggtggcccccctgcccttcgcctgggacatcctgtcccctcagttcatgtacggctccaaggcctacgtgaagcaccccgccgacatccccgactacttgaagctgtccttccccgagggcttcaagtggg<br>agcgcgtgatgaacttcgaggacggcggcgtggtgaccgtgacccaggactcctccctgcaggacggcgagttcatctacaaggtgaagctgcgcggcaccaacttcccctccgacggcccagtaat<br>gcagaagaaaaccatgggctgggaggcctcctccgagcggatgtaccccgaggacggcgccctgaagggcgagatcaagcagaggctgaagctgaaggacggcggccactacgacgctgaggtc<br>aagaccacctacaaggccaagaagcccgtgcagctgcccggcgcctacaacgtcaacatcaagttggacatcacctcccacaacgaggactacaccatcgtggaacagtacgaacgcgccgaggg<br>ccgccactccaccggcggcatggacgagctgtacaagctccacggcctgatcgagtccatgctctgcgatgacactctcatcggcacgcccgagcccgacgagcacccagacccagccatgttcacg<br>gacggcccctgctactccaacggctccgacccgagcagcaccaccacgacgaacccgggcacgcccgtgcagcacgacgacgacctgccgcaggactgcaatcccgagaagggactccggctgctt<br>cacctgctcatggccgccgccgaggcgctctccggcccgcacaagagccgggagctggcacgggtgatattggttcggctcaaggagatggtctccagcaccagcggcaacgctgccgcgtccaaca<br>tggagcgcctcgccgcccacttcaccgacgcgctccaggggctcctcgatgggtcccactccgtcgctgggaccagcaggcaggccgcatcccaccaccacagcaccggcgacgtgttgacggcattc<br>cagatgctccaggacatgtcgccctacatgaagttcggccacttcaccgcgaaccaggcgatcctggaggcggtggcgggcgaccggcgcgtccacatcccaccacgtcctccatggtcac |

| | |
|---|---|
| Extended repair template | ttatataagcgctattgaactttggaacgaacgaaattgctgcacttttcaagtaaacagtttgcggtcattgtggcagctagcgcgcttccgctgccaagatttttcttcttcttgagaacaccttgccaa<br>agtgctacatcagcagaggttagcctttgtagatggtattgggcttatattgggttccatgggcggagtttgtttttcgaacaccagatttataggcatggtaaatcaaatgtttttatggcacatatgtgtt<br>cgctgaggatggcaagtttagttaacagagacggcggccacgacctccacggaagcatgtcaaatttgactcaatttattttttgccaagaaattgtcgtgcttgcaaacttaatttgccaccctgacgc<br>caatataaattgccatacaaaatgtttgatttgccatgcctaatttttttagcatttttgtttcttttatcattagccattatcttcttctttttaaaatcttatagtatgcaaatctaataagattcctggcgagtt<br>attaagaaaaccagattctcattattttttcctttgcaaaaaagagaagattctctctctctcttacaacgattctcattctccggctcaaaaaaaagtttgtttctcattctcttcctgcttaatgcaatcgg<br>tatttttttttttgaggggaacttaatgcaatcagtagagtgcttgcctcgttgctggaaaaagaattgatgatccatgtgatttaacgagaaaaacaaagtccgcccatggtgcccaatattttaggccca<br>gttgggatggtagaacctgctgctggagccagctccagttttgttcgccgaatgccgagtcccggcgcccagggatggctataaataagcgagctcccgtgtccttgtgtacttgtaaaatctgtgctcc<br>ctgcccaccgctctcccctcggttcccacgcgccaaaacattccaacgtggagacaagaagcagcatagcgtgacaacgagggagggagccatggacgtgaccatggaggacgtgatggtgagcaa<br>gggcgaggaggataacatggccatcatcaaggagttcatgcgcttcaaggtgcacatggagggctccgtgaacggccacgagttcgagatcgagggcgagggcgagggccgcccctacgagggca<br>cccagaccgccaagctgaaggtgaccaagggtggcccctgcccttcgcctgggacatcctgtcccctcagttcatgtacggctccaaggcctacgtgaagcaccccgccgacatccccgactacttga<br>agctgtccttccccgagggcttcaagtgggagcgcgtgatgaacttcgaggacggcggcgtggtgaccgtgacccaggactcctccctgcaggacggcgagttcatctacaaggtgaagctgcgcggc<br>accaacttcccctccgacggcccagtaatgcagaagaaaaccatgggctgggaggcctcctccgagcggatgtaccccgaggacggcgccctgaagggcgagatcaagcagaggctgaagctgaa<br>ggacggcggccactacgacgctgaggtcaagaccacctacaaggccaagaagcccgtgcagctgcccggcgcctacaacgtcaacatcaagttggacatcacctcccacaacgaggactacaccat<br>cgtggaacagtacgaacgcgccgagggccgccactccaccggcggcatggacgagctgtacaagctccacggcctgatcgagtccatgctctgcgatgacactctcatcggcacgcccgagcccgac<br>gagcacccagacccagccatgttcacggacggcccctgctactccaacggctccgacccgagcagcaccaccacgacgaacccgggcacgcccgtgcagcacgacgacgacctgccgcaggactgc<br>aatcccgagaagggactccggctgcttcacctgctcatggccgccgccgaggcgctctccggcccgcacaagagccgggagctggcacgggtgatattggttcggctcaaggagatggtctccagca<br>ccagcggcaacgctgccgcgtccaacatggagcgcctcgccgcccacttcaccgacgcgctccaggggctcctcgatgggtcccactccgtcgctgggaccagcaggcaggccgcatcccaccacca<br>cagcaccggcgacgtgttgacggcattccagatgctccaggacatgtcgccctacatgaagttcggccacttcaccgcgaaccaggcgatcctggaggcggtggcgggcgaccggcgcgtccacatc<br>ccaccacgtcctccatggtcactgtgcgtggactacgacctcgccgagggcatccagtgggcgtccctgatgcaggctatgacatcacgacccgatggcgtgtcgcctccgcacctgcgtatcaccgcc<br>atcacgcggagtggcgggggcggcgcgcgggcagtccaggaggccggacggcgcctcgcggccttcgcggggtccatcgggcagcccttctcgttcggacattgccgtctggactcggacgagaggt<br>tccggccggcgaccgtcaggatggtcaagggggagacgctcgtggccaactgcatactccaccaagccgcggcgacgaccaccgtcagacggcccaccggctcggtggcgtccttcttgaccggcat<br>ggcctctctcggggccaaggtggtgacggtggtggaggaggaa |

| | |
|---|---|
| LIR | agtagcaagcagaagcccggcaggtccttagcgaaaaaacggggtgtgctcgcgaactctactctctaccctgcgtgggagtgtgcagaattcacaccgatgggctcggtgtccacggtttaaatattgcaggtttaggtgggaacgcgggaccctgtcttttcggcgcgaaagcgacgggtggtcccgcgtgtggtttgtggctcggggacccgccacgcaggaatctaatattaccctgcgtggcgggtcccgaggcgcactcggcttttcgtgagtgcgccgaggcttttggaccacgtctttatgtcatcacatcaattattggtggtgagtcatcacatattccacctgcaattatgtgccatcgcttagcttataaggaagtgtcggggaaggtatctcg |
| SIR-REP-LIR | acggagtggatgaacacgggtgacggcaagataggcgatattaagaagggtgccctgtatctagtaacctgtactcgtggaggtatcactggagacagtgcctccatttcattcgaagttgtatgtgcctatacgcacgcgtgttacttcaaatccattgggattcaataaataaaatagtattttattcatctcatgtcattcgattacagatgctcggctacgagcaaagataaaccaaactatgacatacaacacactcataaccaaaacatcgaaaaaagaaatacaaggggcgagatcacacaattttagaaaccgtagccgtccgcgctaggacagtcactgcgaagcagtgacattttcgccgaaggcgaagaatgattcaccctcatacatataatgtatcacagcgttagagtacatgtaatccgactgttcaggagtcatatccttgagccaatcttcgtctgggttaactaaaatgatgcaaggtataccaccccgtatcatttttcgcttcccgtacttaggattgacggtgaagtcacgctgagccccgacgaagcacttccagtttggcgtgaacttgaatggaatgtcgtcaattatgttgtacttggcgttgacgtcataggtcgtgaaatcaactaggctgttgtagtagttgtggatccctagagatcttgcccaggaagtctttcctgttcttgtcggaccgcagatgtagatggacttatgccgtcccggtgactcctggaataatcgtccatccactctaaatcagttacggccttatccgcaggagttgaagtacaaaggatatatgattcgaggcttacggagtagagatgttcatttttccagctttcaatggtctcatgacacatgagggactcgatgggaaactcaggtgtgtaagtgctaactgggtctgggaataggtggcgtgcagtgtattcgaagtctttcagacggatagaccattcaaacggaaaacgatggcagaccatgctgagaaattcctctctcgaggtactcgactcaatgatctgtttcatatccgcgtctcggtctttacgacccggagtggtaactgctacgaatgttccccactcagccgtgttgacatcggagtcaacctccttcatgatgtaatcacgaacttggttgcagtctttggcagcttgaatgttaggatgaaaaaatgaaaatggtgatgtgttcataccaatgttgagagcattgggattggtgatggaagcacgaagcttgttttgcacgagtacgtgcagatgtggtgatccatcttcgtggagttccctaactgcagctatgtacagaggttcatatttggccaagagagtgcgaagagagtccaaggcgtactgtggctctaggatgcattgaggatatgttaggaagaggtatttggaatagacacggaacctgggtgcagatgaagaggccatagtagcaagcagaagcccggcaggtccttagcgaaaaaacggggtgtgctcgcgaactctactctctaccctgcgtgggagtgtgcagaattcacaccgatgggctcggtgtccacggtttaaatattgcaggtttaggtgggaacgcgggaccctgtcttttcggcgcgaaagcgacgggtggtcccgcgtgtggtttgtggctcggggacccgccacgcaggaatctaatattaccctgcgtggcgggtcccgaggcgcactcggcttttcgtgagtgcgccgaggcttttggaccacgtctttatgtcatcacatcaattattggtggtgagtcatcacatattccacctgcaattatgtgccatcgcttagcttataaggaagtgtcggggaaggtatctcg |
